# Evaluation of gene expression-based predictors of lymph node metastasis in bladder cancer

**DOI:** 10.1101/2024.11.19.623582

**Authors:** Hafdís Birta Johansson, Fredrik Liedberg, Carina Bernardo, Aymeric Zadoroznyj, Mattias Höglund, Pontus Eriksson, Gottfrid Sjödahl

**Affiliations:** Department of Translational Medicine, Lund University, Malmö, Sweden; Department of Urology, Skåne University Hospital, Malmö, Sweden; Division of Oncology, Department of Clinical Sciences, Lund University, Lund, Sweden

## Abstract

The presence of cancer in pelvic lymph nodes removed during radical surgery for muscle-invasive bladder cancer (MIBC) is a key determinant of patient outcome. It would be beneficial to predict node status preoperatively to individually tailor the use of neoadjuvant chemotherapy and extent of lymph node dissection. Several studies have published node status predictors based on RNA expression signatures in the primary tumor, but none have been successfully validated in subsequent reports.

We use gene expression data and node status from the two largest available MIBC cohorts to evaluate 12 published node-predictive signatures. Additionally, we examine the extent of overlap between differentially expressed genes and signatures across the two datasets, and we train new prediction models which we evaluate in cross-validation and by application to the independent cohort.

All published node status predictors performed either no better than chance or only slightly better than chance in two independent validation datasets (maximal AUC 0.59 and 0.65 and maximum balanced accuracy 0.54 and 0.57 in the two cohorts). Most differentially expressed genes and signatures were only identified in one dataset and only a few, such as upregulation of interferon-response in node negative cases, were enriched in the same direction in both datasets. Transcriptomic predictors trained in one dataset performed poorly when applied to the other independent dataset (AUC 0.60 and 0.62).

In this systematic evaluation, neither the 12 published signatures nor our own models reached an adequate performance for clinical node status prediction in independent data. This indicates that the biological determinants of nodal spread are poorly captured by bulk tumor RNA expression profiles.

## Introduction

Urinary bladder cancer ranks among the leading cancer diagnoses in the Western world. Each year, there are over 613,000 new cases of bladder cancer worldwide and 220,000 deaths, making it the 6th most common cancer in men and the 9th in women [1]. The economic burden per patient is among the highest of all cancer types, attributed to a high risk of disease recurrence, extensive disease surveillance, and costly treatments [2].

For bladder cancer patients, the disease outcome hinges on the stage at diagnosis. Patients with tumors confined to the urothelium (stage Ta) or with invasion only into the lamina propria (stage T1) typically have a favorable prognosis, with only 5-20% progressing to more advanced stages [3]. Conversely, patients with muscle-invasive bladder cancer (MIBC, stage T2-4) face long-term survival rates around 50% despite treatment with neoadjuvant chemotherapy and radical cystectomy (RC). Therefore, there is a need to identify in advance, the patients with micrometastatic disease for whom systemic therapy can make a difference versus patients with disease confined to the bladder who are likely to be cured by radical cystectomy alone. Positive lymph nodes are found in about one in three to one in four patients who undergo radical cystectomy, and they have significantly worse long-term survival after RC compared to node negative patients. [4,5]. The prospect of predicting node status prior to RC has been explored to tailor the extent of lymphadenectomy or to allow the selective administration of neoadjuvant chemotherapy, which is currently standard of care for MIBC patients. [6].

Thus, several studies have used bulk transcriptomic data from tumors to predict node status at RC [7-18]. Although the results in these studies appear promising, none of these published node predictors have been successfully validated in subsequent independent reports, and none of them have reached clinical use.

The aim of this study is to investigate if node predictors perform well enough to be developed into tools for clinical decision-making. To this end, we analyze the expression and predictive performance of 12 published gene signatures in a large cohort of Swedish patients and in The Cancer Genome Atlas (TCGA) cohort. We also compare signature overlap, differentially expressed genes and enriched signatures in the two cohorts to identify any robust biological basis for metastatic lymph node dissemination in bulk expression data. Finally, we estimate to what extent node status can be predicted from bulk RNA expression data by performing nested feature selection followed by cross-evaluation of new classification models derived in each of the two cohorts.

## Materials and methods

### Identification of published lymph node status prediction signatures

We screened the literature for articles related to classification and/or prediction signatures for lymph node metastasis in bladder cancer. Until November 1^st^, 2023, we identified 12 studies, published between 2006 and 2023 that involved tumor RNA expression and node status prediction (Table 1). The study by Mitra et al. (2014) [10] was included despite being developed as a post-cystectomy recurrence signature, because the authors used it to predict node status in subsequent work [11]. We also included Stubendorff et al. [14] despite being developed from methylation data, because the study reported correlation between methylation and expression. Half of the identified studies utilized data from The Cancer Genome Atlas bladder cancer (TCGA-BLCA) cohort, which included a mix of biospecimens collected from preoperative transurethral resection and radical cystectomy tumor tissue. The remaining six studies were exclusively based on tissue analyses from radical cystectomy specimens.

**Table 1.**
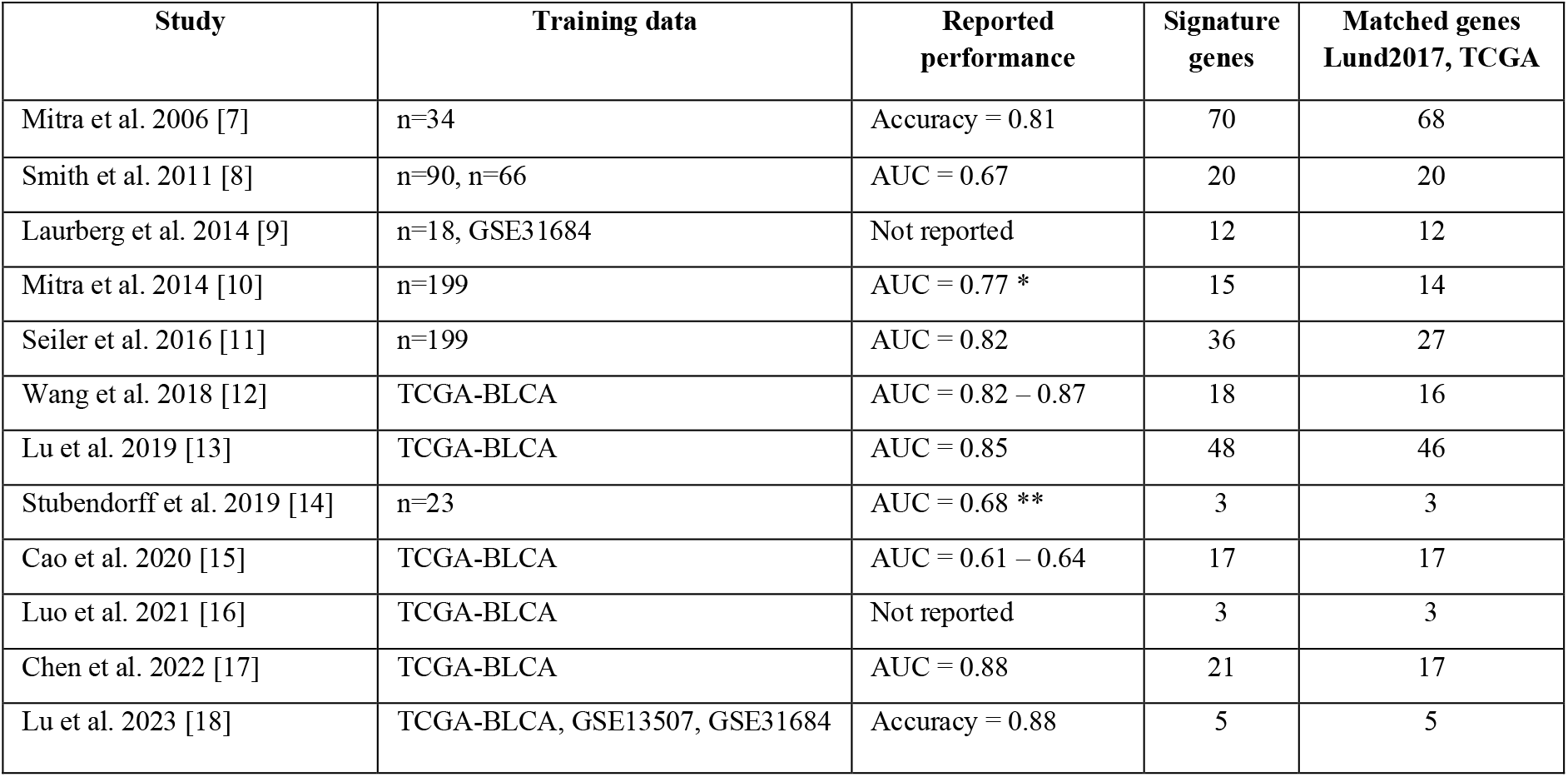
Identified lymph node status predictive signatures. * Mitra et al. 2014 developed a post-cystectomy recurrence risk predictor, and the AUC is for that outcome. ** Stubendorff et al. 2019 developed a methylation panel and the AUC is for that data type.

| Study | Training data | Reported performance | Signature genes | Matched genes Lund2017, TCGA |
| --- | --- | --- | --- | --- |
| Mitra et al. 2006 [7] | n=34 | Accuracy = 0.81 | 70 | 68 |
| Smith et al. 2011 [8] | n=90, n=66 | AUC = 0.67 | 20 | 20 |
| Laurberg et al. 2014 [9] | n=18, GSE31684 | Not reported | 12 | 12 |
| Mitra et al. 2014 [10] | n=199 | AUC = 0.77 * | 15 | 14 |
| Seiler et al. 2016 [11] | n=199 | AUC = 0.82 | 36 | 27 |
| Wang et al. 2018 [12] | TCGA-BLCA | AUC = 0.82 – 0.87 | 18 | 16 |
| Lu et al. 2019 [13] | TCGA-BLCA | AUC = 0.85 | 48 | 46 |
| Stubendorff et al. 2019 [14] | n=23 | AUC = 0.68 ** | 3 | 3 |
| Cao et al. 2020 [15] | TCGA-BLCA | AUC = 0.61 – 0.64 | 17 | 17 |
| Luo et al. 2021 [16] | TCGA-BLCA | Not reported | 3 | 3 |
| Chen et al. 2022 [17] | TCGA-BLCA | AUC = 0.88 | 21 | 17 |
| Lu et al. 2023 [18] | TCGA-BLCA, GSE13507, GSE31684 | Accuracy = 0.88 | 5 | 5 |

### Validation datasets

The signatures were evaluated in the Lund2017 and the TCGA cohorts. Lund2017 is a consecutive cohort of 307 patients who underwent radical cystectomy in southern Sweden between 2004 and 2011 [19]. Of the 307 patients, pathologic node status was available for 296, of which the majority (92%) underwent an extended lymphadenectomy. The utilization of neoadjuvant systemic therapy in this cohort was low (9.4 %) [20]. RNA expression data (Affymetrix Gene ST 1.0) is available at the Gene Expression Omnibus data repository under accession number GSE83586. For this study, probe set data following RMA and ComBat adjustments for different labeling batches was annotated with the BrainArray V25 based on Gencode 36, summarized at the gene level, and median centered. This resulted in data from 22383 genes with a unique HGNC gene symbol. RNA-seq data from The Cancer Genome Atlas (TCGA) urothelial bladder cancer cohort was obtained through the TCGABiolinks R package, normalized using TMM (Trimmed mean of M values) into log2+1 values and median centered. Ensembl gene ID was matched to the Lund2017 cohort including the full set of 22383 genes in both validation datasets. The full datasets were used to extract genes from published signatures. Metadata including node status for 364 TCGA tumors was obtained from the original report [21]. Two patients were excluded due to inconsistent node status in different versions of the metadata. Filtered datasets excluded genes with 0 values in more than 50% of cases before normalization. The resulting sets containing 16297 genes were used for differential expression analysis and to identify novel node-predictive signatures.

### Statistical analyses

Fisher’s exact test was used to test the association between categorical variables and the difference between observed and expected overlap of gene lists. The association between continuous variables (e.g. signature scores) and node status were analyzed by receiver operator characteristic (ROC) curves and calculating the area under the ROC curves (AUC) using ‘ROCR’ and ‘ggROC’ R packages. Cancer-specific and overall survival for the Lund2017 and TCGA cohorts, respectively, were compared by log-rank test using the ‘survival’ R package.

### Molecular classification, differential expression, and gene enrichment analyses

Molecular subtype classification of both cohorts according to the Lund taxonomy was performed using the ‘LundTax2023Classifier’ R package [22]. Differential gene expression analysis comparing node positive versus node negative cases was performed using the R package ‘limma’ with Benjamini-Hochberg corrected p-value ≤ 0.05 considered as significant. Enrichment analysis of gene ontology (GO) terms was performed using Panther [23], with the annotation dataset: GO biological process complete. The 5% most differentially expressed genes were tested against the full set of 16297 genes used as reference background. Pre-ranked gene set enrichment analysis (GSEA software v4.3.2 [24]) was performed differential expression analysis, scanning for the following MSigDB gene sets: H (hallmark), C2 (curated), C6 (oncogenic).

### Mean signature scores

The features/genes from each published node predictor were translated to the corresponding gene symbol and extracted from the Lund2017 and TCGA datasets. Then, a *mean signature score* was defined as the mean of the normalized and median centered log2 expression values for the signature genes in each dataset. If the signature consisted of both up- and down-regulated genes in node positive patients, the ratio of mean scores (Mean log2-Exp up – Mean log2-Exp down) was used. Each gene’s directionality was obtained from the original report when the information was available. When individual genes’ directionality could not be obtained from the original report, it was inferred from the expression in the dataset, i.e. it was set to be up-regulated in node positive if the gene’s expression in node positive cases was above its median expression value in the dataset.

### Node status prediction models

In addition to mean signature scores, we used logistic regression and random forest (RF) models to explore the external signatures’ potential to predict node status. The ‘glmnet’, ‘nestedcv’, and ‘caret’ R packages were used to perfom 5 fold nested cross-validation of each signature and to build a final binomial classification model (5 outer folds, using 5 inner folds assessing hyperparameter α spanning 0 to 1 in 0.05 increments). This was repeated 10 times for each signature, using predefined training and test folds, to estimate each signature’s performance in 50 outer cross-validation folds. Signature evaluation through RF was performed with the ‘ranger’ and ‘caret’ R packages using each signature’s genes as features in a 5-fold cross validation approach, repeated 5 times, with performance assessed on the 25 outer cross-validation folds. Final models were applied both to the dataset in which it was trained and to the fully independent data.

The same logistic regression and RF approach was used to evaluate and test the performance of the top 50 differentially expressed genes identified in the Lund2017 and TCGA datasets. Finally, the Lund2017 and TCGA datasets were used to evaluate the performance of logistic regression models that utilized fully nested feature selection to avoid information leakage between training and test folds. Four feature selection methods; glmnet, t-test, Wilcoxon test, and ranger, were used to identify top features in each training fold for each dataset. The internal performance of this approach was evaluated on the 50 outer folds from 5-fold nested cross-validation repeated 10 times. The full models were applied to both datasets to contrast with the cross-validation performance metrics.

## Results

### Identification of node status prediction signatures and validation datasets

We identified 12 published gene-expression based node predictors by literature review [7-18] (Table 1). We used the Lund2017 cohort (n=296) and the TCGA cohort (n=362) as validation datasets with 33% and 35% of patients being node positive, respectively. Based on the 12 published gene signatures and these two validation cohorts we devised a strategy to analyze node status prediction in three steps outlined in Figure 1A. In the Lund2017 cohort, the median number of removed lymph nodes was 28 for node negative patients and 29 for node positive patients (interquartile ranges (IQRs) 18-39 and 20-39, respectively). In the TCGA cohort both node negative and node positive patients had a median of 18 removed lymph nodes (IQRs 9 – 28 and 12 – 37, respectively). The distribution of RNA-based molecular subtypes according to the Lund taxonomy did not differ between the cohorts (p >.05, Fisher’s exact test). We observed an enrichment of node positive cases in the ‘Genomically unstable’ subtype in the TCGA cohort (26 of 44 cases of this subtype (59%) were node positive, p=.009, Fisher’s exact test). However, this finding was not observed in the Lund2017 cohort (p=.731) (Figure 1B). We then confirmed the prognostic nature of pathologic node status by univariable survival analysis in the Lund2017 and TCGA cohorts (Hazard ratios 3.7 (95% CI 2.5 – 5.4) and 2.1 (95% CI 1.5 – 2.8) (Figure 1C).

**Figure 1.**
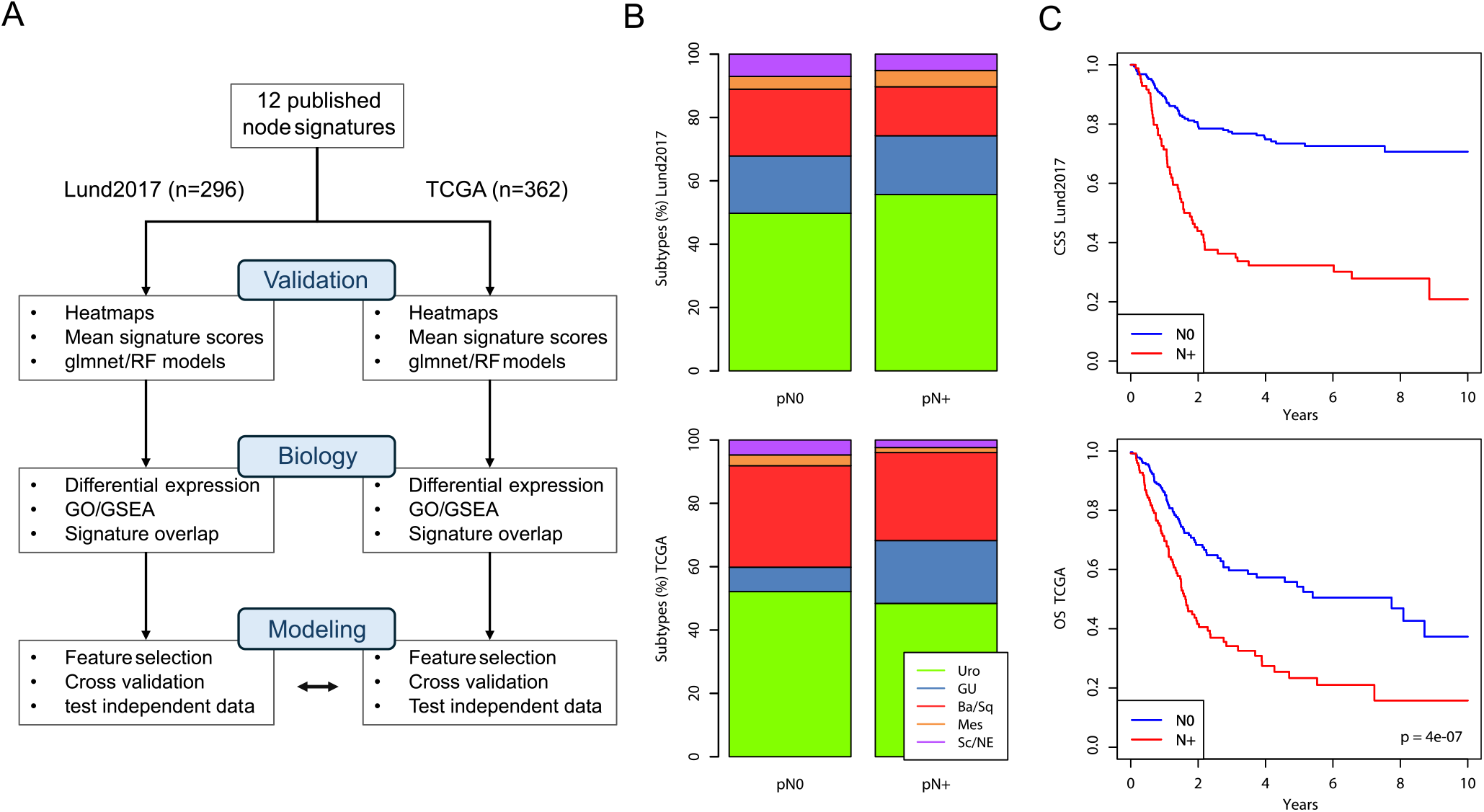
Public datasets and workflow for evaluating node status prediction in UC. A) Schematic outline of the workflow in this study. In the first step, 12 gene expression signatures from the literature were extracted from the Lund2017 and TCGA cohorts and validated using mean signature scores as well as glmnet and RF models. In the second step, the biological underpinnings of node status were analyzed in the two cohorts. In the third step prediction models were trained and evaluated by cross-validation, and the final models cross applied to the other dataset. B) Association between molecular subtype (LundTax classification system) and pathologic node status in the Lund2017 and TCGA cohorts. C) Cancer-specific, and overall survival stratified by dataset in the Lund2017 and TCGA cohorts, respectively.

### Evaluation of published node-predictors

We extracted 12 published gene signatures from the two validation datasets and visualized gene-expression as heatmaps ordered first by dataset and then by molecular subtype (Figure 2). Visual examination of the heatmaps identified no genes obviously associated with node status, whereas several signatures appeared largely linked to molecular subtype biology (Smith11, Mitra14, Chen22). Other signatures (Laurberg14, Cao20, Luo21) identify stromal components with high expression in the Mesenchymal-like and Basal/Squamous subtypes.

**Figure 2.**
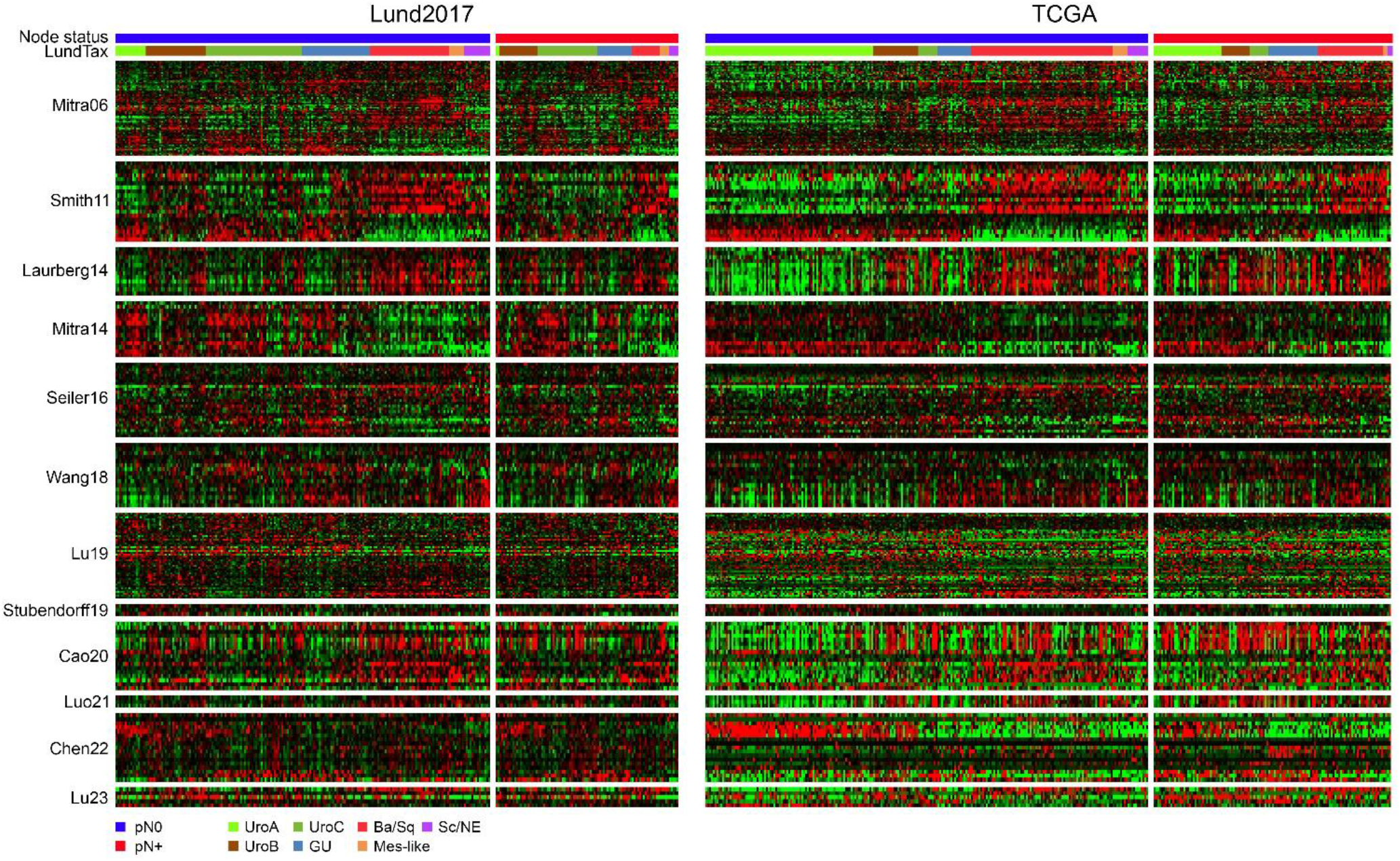
Expression of published node-signature genes in the Lund2017 and TCGA cohorts. A) Heatmaps showing the expression of 12 published gene signatures in the Lund2017 dataset arranged by node status and molecular subtype. Genes within each signature were ordered by hierarchical clustering. B) Heatmaps showing the same genes in the same order as in A) in the TCGA dataset arranged by node status and molecular subtype.

Among the 12 signatures, there were 6 overlapping genes, compared to the 1.2 expected by chance (Figure 3A). However, 5 of the 6 overlapping genes were found between studies derived from the TCGA dataset and only one gene overlapped among studies with independent training datasets. We calculated mean signature scores and derived ROC-curves and AUC values for lymph node metastasis in both datasets. In the Lund2017 data, the Mitra06 mean signature score performed slightly better than the other signatures with an AUC of 0.63 (Figure 3B). Of the 12 signatures, 6 were originally derived from the TCGA data and could not be independently tested in Figure 3C (included as dashed lines). The best performing mean signature in TCGA data, excluding those derived from this dataset, was Mitra06 with a moderate AUC of 0.68, surpassed only by four TCGA derived signatures that reached AUCs of 0.69-0.73.

**Figure 3.**
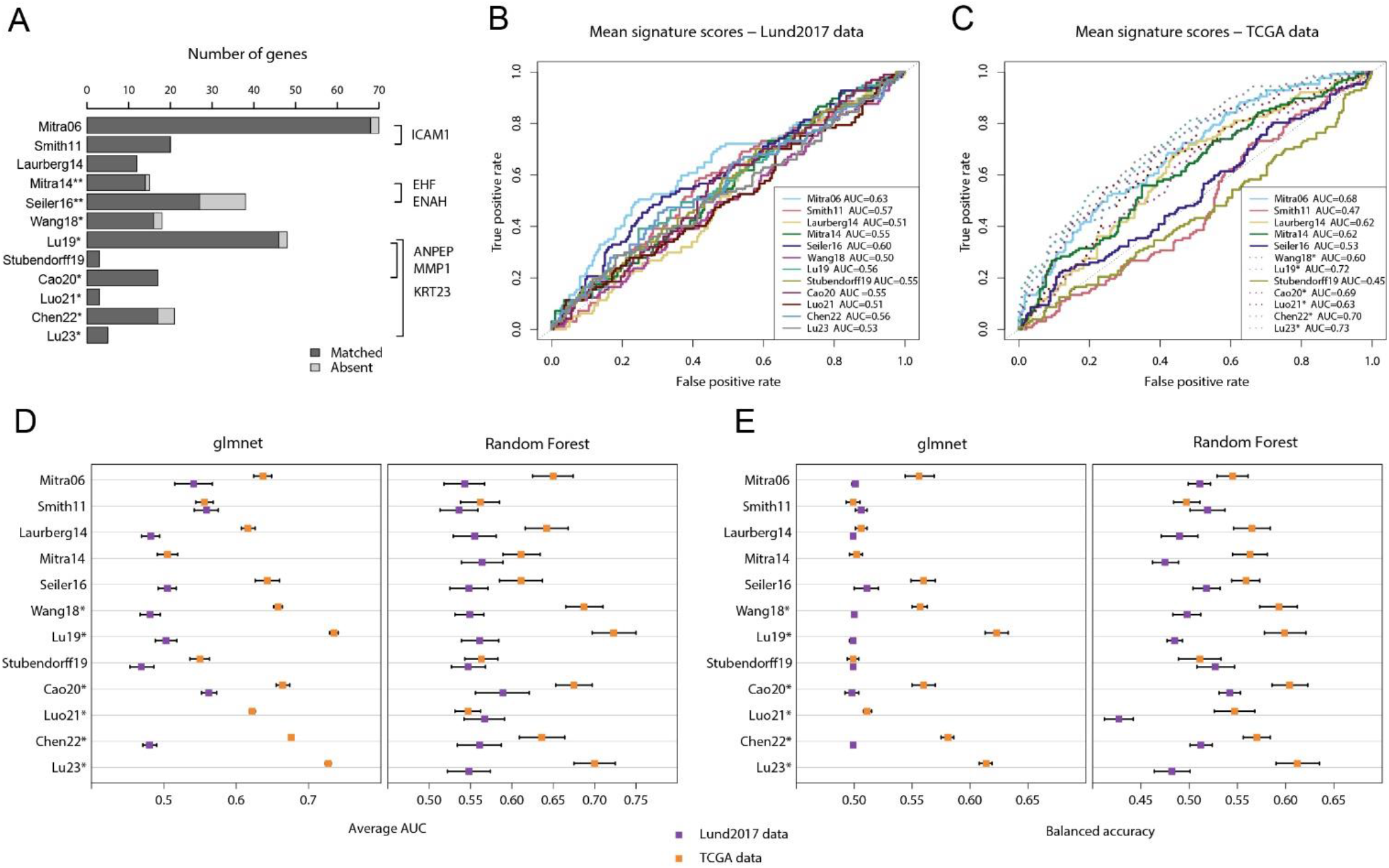
Published node-signatures do not stratify or independently predict node status in the Lund2017 and TCGA cohorts. A) Barplot showing number of genes and gene overlap between the 12 published node-signatures. Five out of six overlapping genes were from studies with the same training dataset (studies marked with * or **). Thus, only *ICAM1* was independently identified in two signatures. The expected number of overlapping genes based on simulation was 1.2. B) ROC-curves and AUC-values are shown for the mean signature scores of the 12 published signatures in the Lund2017 dataset and in C) the TCGA dataset. Signatures derived from the same dataset in C) are marked with an asterisk and plotted with dotted lines. Note that gene directionality in B) and C), when not provided in the original study, was inferred based on expression in node positive versus node negative tumors dataset, possibly leading to inflated AUC-values. D) Signatures’ performance is shown as average AUC-values in repeated nested validation using glmnet and RF models trained on the Lund2017 or TCGA datasets. E) Signatures’ performance is shown as balanced accuracy in repeated nested validation using glmnet and random forest (RF) models trained on the Lund2017 and TCGA datasets. Results from glmnet models based on the Mitra14, Luo21, and Lu23 signatures in the Lund2017 dataset are not shown because all variables were rejected.

In addition to these simple mean signature scores, we constructed models using logistic regression (glmnet) and random forest (RF). The signature that performed best with glmnet and RF models in the Lund2017 data was Cao20, with average AUC of 0.56-0.59 and balance accuracy of 0.50-0.54 (Figure 3D-E). Results in the TCGA dataset were only slightly better, with the best performing non-TCGA derived signatures being Seiler16 and Mitra06, obtaining average AUC-values of 0.61-0.65 and average balanced accuracy of 0.55-0.56 in cross-validation (Figure 3D-E).

### Limited shared biology pertaining to node status in the Lund2017 and TCGA cohorts

Next, we performed parallel differential expression analysis according to node status and subsequent gene set enrichment analysis (GSEA) in the two datasets to test if they would arrive at the same conclusions. Differentially expressed genes (DEGs) in the two cohorts are shown in Figure 4A. After adjusting for multiple testing, one gene (*HSPA5*) was significantly associated with node status in the Lund2017 dataset, whereas 910 genes were significant in the TCGA dataset. To assess if the two cohorts showed similar trends in differential gene expression, we compared the observed and expected overlap among the top 0.1%, 1%, 5%, and 10% of up- and down-regulated genes, regardless of statistical significance (Figure 4B). Among the 10% most up- and 10% most down-regulated genes in Lund2017 and TCGA data, 225 and 257 genes overlapped compared to 163 expected by chance alone (both p<.001). At the 5% level, 70 and 79 genes overlapped, compared to 41 expected by chance (p<.01, p<.001). For the 1% top up- and down-regulated genes in node positive patients 5 genes overlapped (*COMP, ELN, FAT3, PCP4, PRELP*), while 1.6 was expected (p>.2). We then performed gene ontology (GO) term and Gene set enrichment analysis (GSEA). The most overrepresented GO terms in node positive tumors were related to muscle-cell function in Lund2017 and collagen, elastic fiber, and extracellular matrix organization in TCGA. In the node negative tumors, the most overrepresented GO terms in Lund2017 involved antigen processing, type I interferon signaling, antigen presentation via MHC class I, and endoplasmic reticulum to Golgi transport. In the TCGA dataset several terms related to RNA processing, splicing and translation were overrepresented in node negative cases (Supplementary Table 1). The only overrepresented GO terms with the same directionality in the two cohorts were high-level terms related to RNA processing in node negative cases. We also performed GSEA using the MsigDB Hallmark, Curated, and Oncogenic gene set categories. For both datasets, a large number of signatures were found to be significantly enriched in either nodal category, but only a small fraction of them in both datasets (up in node positive: 22 of 729, 3%; up in node negative: 78 of 1834, 4%) (Supplementary Table 2) (Figure 4C). Among the shared signatures enriched in node positive tumors were six signatures of genes with CpG-rich promoters that have been found to be epigenetically silenced upon reprogramming to pluripotent stem cells. Node positive cases in both datasets were also enriched for a muscle/myosin signature derived from head and neck squamous cell carcinoma and a breast cancer chromosome 1q21 amplification signature. On the DNA level the 1q21-22 amplified cases in TCGA had larger proportion with node positive disease compared to cases without 1q amplification (Fisher’s exact test, p=0.0021) (Figure 4D). The node negative cases in both datasets were enriched for hallmark MYC targets signatures as well as mRNA processing/splicing and translation. Furthermore, signatures for interferon signaling, viral infection, and antigen presentation via MHC class I were highly enriched in node negative cases in the Lund2017 dataset but only moderately so in TCGA data. We tested an interferon gamma response score defined in urothelial cells [25], but this score was not significantly associated with node status in either dataset (Lund p=0.16, TCGA p=0.96, Supplementary Figure 1). While the observed commonalities suggest that there are shared processes linked to the propensity for lymphatic dissemination, the overlapping results were in minority and, in general, the findings correlated with node status in one cohort did not show the same pattern in the other cohort.

**Figure 4.**
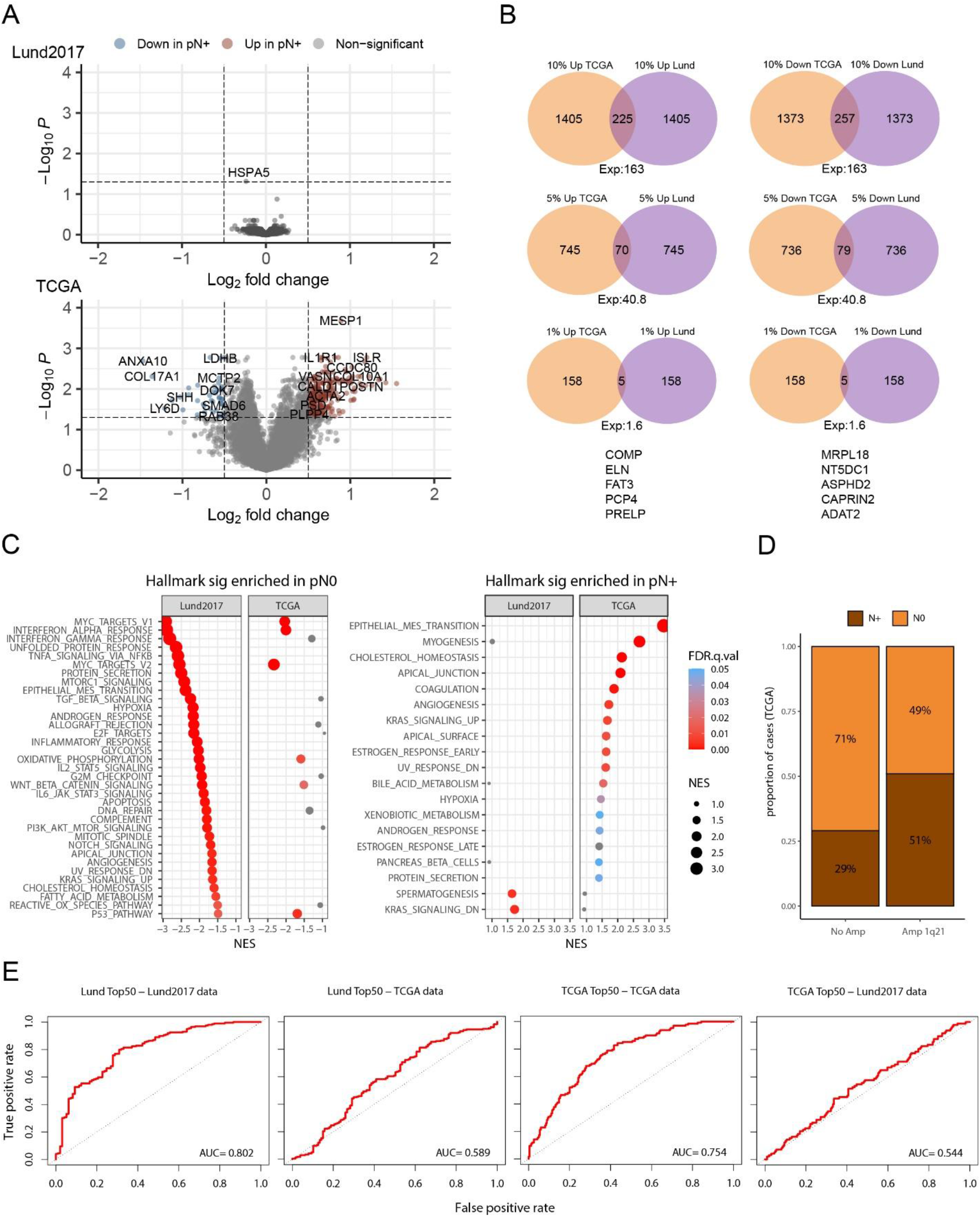
Differential expression analysis by node status in the Lund2017 and TCGA cohorts. A) Volcano plot showing differentially expressed genes in the Lund2017 and TCGA datasets. In Lund2017, the only significant gene, HSPA5, had modest Log2 fold change of -0.24. Highlighted genes in TCGA show absolute Log2 fold change > 0.5 and adjusted p-value < 0.05. Venn diagrams in B) show the overlap between lists of up- and downregulated genes of various sizes (10%, 5%, 1% and 0,1% of all genes) in Lund2017 and TCGA along with the expected number of overlapping genes for random gene lists. C) Hallmark mSigDB gene signatures with positive enrichment in pre-ranked GSEA for upregulation in node negative (pN0) and node positive (pN+) cases. Only a minority of significant (FDR <0.05) signatures maintained directionality and significance in both datasets. D) Proportion of node positive cases was higher among tumors with 1q21-22 amplification in the TCGA cohort (p=0.0021, Fisher’s exact test). E) ROC curves showing the performance of mean signature scores based on the top 50 DEGs (25 up- and 25 downregulated genes) in Lund2017 and TCGA to separate node status in the two datasets. High AUC values were achieved when signatures were tested on the same dataset in which feature selection was performed, whereas AUC values were low when cross-applied to the other dataset.

We then asked if the DEGs in Lund2017 could be used to predict node status in TCGA and vice versa. First, the top 25 up- and down-regulated genes in each dataset were used to calculate mean signature scores as well as glmnet and random forest (RF) prediction models in both datasets. The mean signature scores from the DEGs could only predict node status in the cohort from which the DEGs were defined (Figure 4E). The more advanced glmnet and RF methods also yielded comparable results, namely good performance in cross-validation in the cohort in which the selected genes were differentially expressed (average AUC 0.76-0.80 and average balanced accuracy 0.63-0.70). However, the performance in cross-validation was poor for all models constructed using the other cohort’s DEGs (average AUC and balanced accuracy of 0.62-0.67 and 0.56-0.57 for models trained in TCGA on Lund2017 DEGS, and 0.56-0.58 and 0.50-0.53 for models trained in Lund2017 on TCGA DEGs) (Supplementary Table 3). This indicates that model performances are overestimated when evaluated in cross validation, or in a left out “test set”, when the initial feature selection is performed on the same full training data.

### Developing and cross-evaluating new node-predictors in the Lund2017 and TCGA cohorts

Finally, we performed repeated nested cross-validation of glmnet logistic regression models using internal feature selection only on the current training fold. This insulates both hyperparameter optimization and feature selection from the testing folds to prevent information leakage. Feature selection was performed either by glmnet alone, or by first identifying the top 100 genes through t-test, Wilcoxon test, or RF variable importance. The maximum average AUC from this cross-validation approach was 0.57 for Lund2017 and 0.67 for TCGA (Figure 5A).

**Figure 5.**
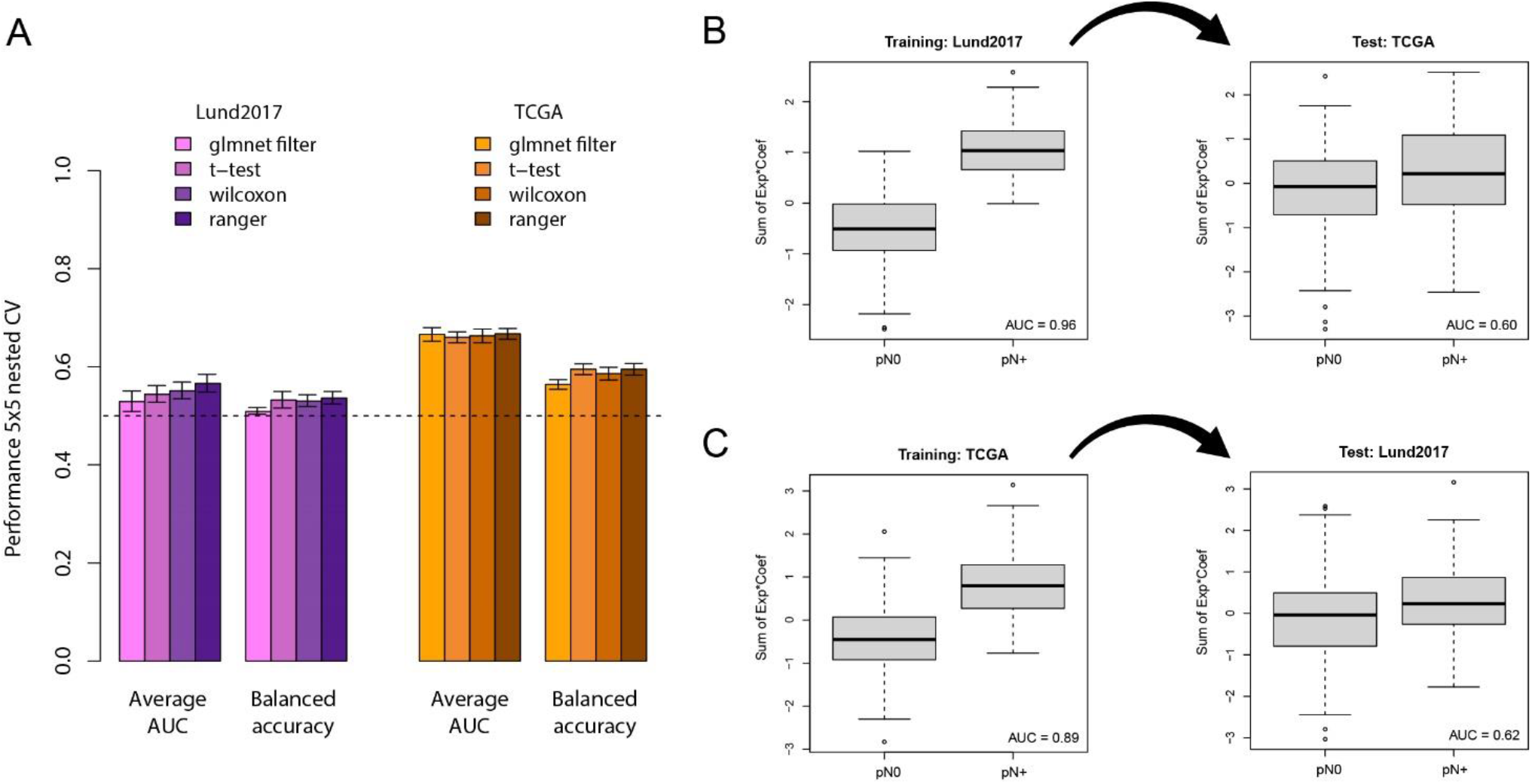
Training and cross application of node status classification models in the Lund2017 and TCGA cohorts. A) Barplots showing the average performance (AUC and balanced accuracy) across repeated cross-validations in the two datasets using glmnet models with four different feature selection methods. B) The glmnet model with the ranger feature selection method has a nearly perfect separation of node status in the training data (fitted to the Lund2017 dataset), but fails when applied to the independent TCGA dataset. C) The glmnet model with the ranger feature selection method trained on the TCGA dataset also shows good separation of node status in the training dataset it was fitted on and not when applied to the independent Lund2017 dataset.

We then contrasted these cross-validation results with those obtained when feature selection is performed on the full cohorts. For both Lund2017 (Figure 5B) and TCGA (Figure 5C), performing feature selection on the whole dataset resulted in very high AUCs in the training data (0.96 and 0.89, respectively). However, lower AUCs of 0.60 and 0.62, similar to those obtained in nested cross-validation with internal feature selection, were obtained when the models were cross-applied to the independent data. Thus, independent validation results and adequate cross-validation estimates without information leakage both show transcriptomic node-predictors perform worse than what would be required for a useful clinical prediction tool.

## Discussion

Current neoadjuvant cisplatin-based regimes give rise to significant adverse effects, while the absolute therapeutic survival benefit for bladder cancer patients is only about 5-10% [26,27]. Therefore, a key aim of the research field is to restrict the use of systemic therapy to patients for whom it is most likely to improve outcomes. Some patients who undergo radical cystectomy and pelvic lymph node dissection can be cured despite having positive lymph nodes [28], but most node positive patients will experience disease recurrence. On the other hand, patients who are node negative at cystectomy are more likely to have a disease limited to the bladder and thus have a greater chance of cure by cystectomy alone [4,5]. Retrospective studies have found a clinical benefit of extended versus limited lymph node dissection [29], but the two randomized studies on the topic have both been negative [30,31]. Hence, an accurate prediction of node positivity in clinically node-negative patients before radical cystectomy has potential to identify both patients who are more likely to benefit from neoadjuvant chemotherapy, and patients for whom a less extensive lymph node dissection could be beneficial.

Several studies have constructed node status prediction models based on data available before cystectomy, including clinical parameters [32-35] and by applying deep learning to histology slides [36,37]. The first RNA based node prediction study we identified was published in 2006, and since then almost one such study per year has been published. No meta-analyses or systematic independent validation studies have been performed that compare all the signatures. In our initial analysis of mean signature scores, the only signature that potentially performed better than chance in both datasets was Mitra06 [7]. However, the mean signature approach may have overestimated the signatures’ true value since each gene’s directionality had to be inferred from the test data when not specified in the original report. A potential reason why the Mitra06 signature showed the best performance could be that it included the largest number of genes, thereby gaining more advantage from the inferred directionality. Here it is important to note that the study of Mitra et al. [7] emphasizes that the predictive value lies in the relationship between gene’s expression level and not in their absolute values. Importantly, we therefore also evaluated each signature’s predictive information content using glmnet and random forest models. The best performing signatures in Lund2017 and TCGA showed cross validation AUC-values of 0.59, and 0.65 and balanced accuracies of 0.54 and 0.57, respectively. Taken together, our methods show that all 12 signatures performed no better, or minimally better than chance in these two cohorts. Six of the signatures that were derived in the TCGA dataset performed, as expected due to overfitting, better in TCGA data, but they performed as poorly as the non TCGA derived signatures in the Lund2017 dataset. Despite the intended use on trans-urethral resection of bladder tumor (TURBT) specimens, all signatures were developed on cohorts that, at least partly, used tissue specimens from RC. The source of tissue specimen may have systematic effects on the tumor expression profile, and only one study [8] addressed this potential bias.

There was only one overlapping gene between studies with independent training data (*ICAM1* in Mitra06 and Smith11), which was no greater than the overlap expected by chance. Few previous validation studies of these node-predictors exist in the literature. Notably, van Kessel and colleagues [38] performed a thorough qPCR validation-study of the Smith11 signature [8] and found that the signature had no predictive value. Seiler and colleagues, when testing their own signature, also attempted to validate the Mitra14 signature [10], again without success [11]. Similarly, Wang and colleagues compared their own results to those obtained with the Smith11 [8], Laurberg [9], and Mitra14 [10] signatures. Curiously, this validation in two external datasets resulted in very high AUC-values (0.74-0.92 and 0.81-0.83) for all four signatures [12]. Notably, one dataset used for validation by Wang and colleagues (GSE13507) contains 63% Ta-T1 tumors. Such early-stage tumors are not candidates for cystectomy, have no pathologically determined node status, and cannot serve to validate this type of predictor. The description of how the signatures were applied to the validation datasets did not allow us to reproduce the findings by Wang et al. We note that the same GSE13507 dataset which contains many Ta tumors was also used by Lu et al. for model training [18]. In our opinion, this undermines the foundation of the Lu23 signature. In summary, most previous validation attempts have been negative, and some studies have clear methodological flaws.

We then asked if the absence of predictive information in the published signatures was because each study had failed to identify the correct node-predictive genes, or because no node-predictive genes exist in the bulk tumor transcriptome. To answer this question, we performed differential expression analysis and constructed prediction models in both datasets to see if one would arrive at similar conclusions. Top differentially expressed genes showed more overlap than expected by chance, but the observed versus expected overlap was modestly increased and most overlapping genes were false positives resulting from random chance. Nearly all GO-terms and GSEA results were dataset specific, with a few exceptions: A breast cancer signature of 38 genes located on chromosome 1q21 was enriched among node positive patients in both datasets. This amplification is common in bladder cancer and the association with node status was confirmed on the DNA level. Among the node negative patients, the shared signatures of interest included type I interferon response, antigen presentation, and MYC targets. For the first two, a hypothetical explanation is that tumors may solve the adaptive immunity problem either by editing or by escape [39]. Tumors that evade destruction by immunomodulation would exhibit higher expression of interferons and antigen presentation proteins and would also be inefficient at colonizing an immune-rich niche such as the lymph nodes, outside of their immune-modulated environment in the bladder wall. These hypotheses merit further investigation, but the modest normalized enrichment scores in the order of 2-3 indicate that the underlying processes only contribute marginally to the phenomenon of lymphatic dissemination. The finding that top differentially expressed genes in one dataset were poorly predictive of node status in the other dataset (max AUC 0.60 – 0.62) further supports that any links that may exist between the tumor transcriptome and node status will be multi-factorial or context dependent.

The metastatic process includes many steps which are reflected in the expression profiles of cells within the tumor [40], so how come node status is poorly predicted from bulk RNA expression data? It is possible that a temporal or spatial resolution is needed to capture transient or local biological processes governing metastatic spread [41]. It could also be that direct measuring of other processes relevant to node-metastasis, e.g. epigenetic states [42,43] or immune evasion [44] would better identify tumors in which nodal spread has likely occurred. It is also a possibility that some process(es) involved in lymph node metastasis are underdetermined or stochastic. In that case, it would be difficult to predict metastatic status based on the primary tumor even with access to spatiotemporally resolved multi-omic data.

This work has many limitations. We chose to only analyze gene expression signatures, a subset of node status predictors in bladder cancer. Furthermore, since most of the studies did not provide sufficient information to enable external application of prediction models, we did not apply the published models as they were originally created. Instead, we evaluated the signatures’ information content by multiple other methods including training of new prediction models based on the signature genes. An important implication of our results is that caution is warranted when evaluating node status prediction models in a single dataset. The feature filtering/selection step, in particular, should not be influenced by the cases that are subsequently used for model evaluation. Doing so will overestimate the performance even if separate test sets or out-of-bag samples are used to set model parameters. Another benefit of our work is that any new study in this field can be assessed in Lund2017 and TCGA (node status available in Supplementary Table 4) and be directly compared to our results before publication.

The poor result of all 12 bulk transcriptomic signatures and of our own prediction models trained and cross-applied on two large cohorts suggests that new or alternative approaches will be needed to identify clinically useful node status predictors.

## Acknowledgments

We thank Benjamin Ulfenborg at University of Skövde for helpful discussions around the evaluation of prediction models.

