## Supplementary figures and images for "Evaluation of gene expression-based predictors of lymph node metastasis in bladder cancer"

### Supplementary Figure 1

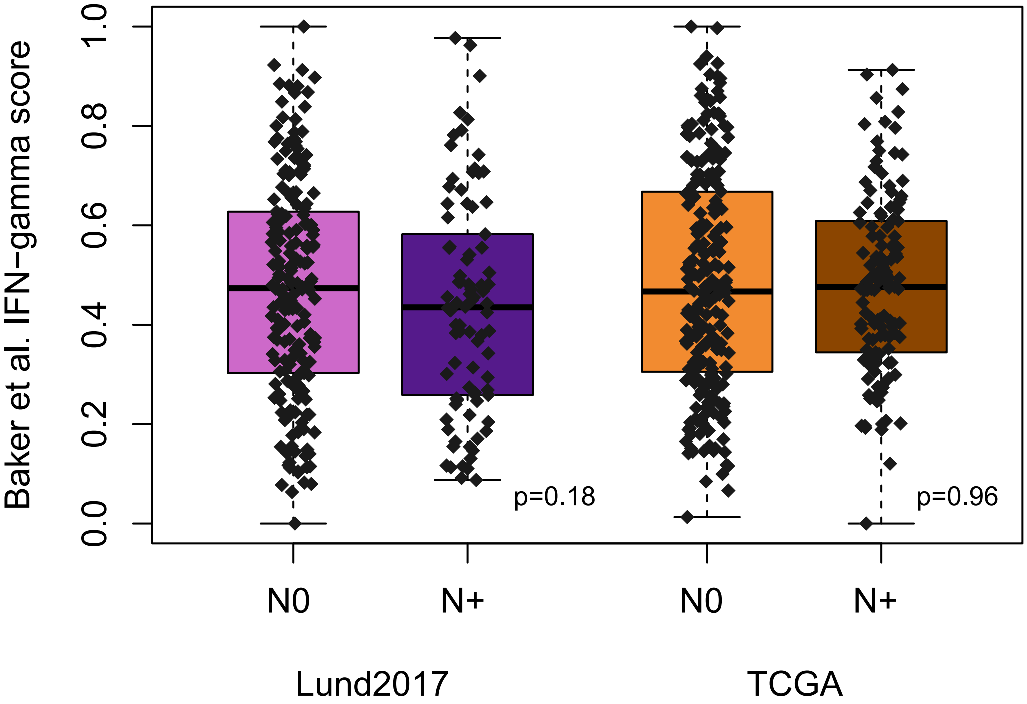
